# Partial illustration of human sperm DNA via microscopy and quantitative analysis of nucleotides

**DOI:** 10.1101/2022.02.18.481020

**Authors:** Jaleh Barzideh

**Affiliations:** Infertility Research Center, Shiraz University of Medical Sciences, Shiraz, Iran; HMRI/Life Science School, University of Newcastle, Newcastle, NSW, Australia

**Keywords:** nucleotides average, 5-methylcytosine, sperm chromatids, CMA3 staining

## Abstract

**Background:** general structure of human sperm has not been profiled yet. Human sperm DNA characterization should progress the medical diagnostic and therapeutic methods rather than developing biological sciences. The aim of the present study was to provide biological insights into the common structure of human sperm. The value of this investigation is establishing an initial basic map of sperm head structure that leads to further advanced standardization of normality in this creature. For this purpose, analytical and microscopic methods were applied.

**Methods:** High-performance Liquid Chromatography (HPLC) and flow cytometry were hired to quantify the DNA compositions. As well fluorescent, confocal and advanced light microscopy was applied to identify the stained sperm DNA by chromomycinA3 (CMA3) and 5-methylcytosine antibody (5-mc)

**Results:** HPLC demonstrated the mean values of nucleotide bases’ percentage in the structure of the sperm DNA regardless of the fraction that sperm was collected from gradient wash, sequenced from 27.6%, 8.92%, 27.05% and 35.36%. Also, quantitative flow cytometry of global 5-methylcytosine showed not a regular fluctuation in individuals with normal sperm while, there is a permanent increase in 50% fraction collected from percoll gradients.

CMA3-positivity levels as well, were negatively correlated with sperm quality harvest by percoll gradients (p<0.0001), and positively correlated (*P*<0.05) with global methylation as determined by flow cytometry. Interestingly, in this text microscopy of immunocytochemistry of sperm cells stained by CMA3, demonstrated a different view from cells’ heads.

**Conclusions:** obviously these explorations suggest some new possibilities in assessment of rough chemical level of nucleotides and cytochemistry of sperm head structure. The chromatin brightness presented with CMA3 by microscopy shows a direct relation with more extensive DNA methylation in sperms collected from low gradients of percoll wash. While, fluctuated 5-methylcytosine levels show personal presentation and even exclusive to individual sperm expression. This study induces further research on new assumptions in nuclear equilibrium in the axiom of DNA ladder in related to 5-mcytosine level in human sperm.

## Background

Understanding chromosomal morphology of human sperm and molecular cell biology might cause a leap in diagnostic methods and further advanced therapeutic cell manipulation. Robust sophisticated techniques and analysis have been used for estimation of the structure of chromatids and geometry of DNA in spermatozoa. In the current text, both novel results of microscopy and analytical methods could enhance future biomed technology and modeling by revealing micrographs of chromosomes and the micro components of the sperm structure.

Recent attentions attracted to detail of structural characters of sperm nucleus ^1^ and intra cellular localization of chromosomes^2^. Physical characters of this cell such as telomere lengths used as a fertility diagnostic sign ^3^. Which might be related to the importance of telomeres as the first point of paternal DNA that would be in contact with the ooplasm ^4^. Different theories have been developed about the location of chromosomes and centromeres as well as its tendency to clustering in sperm heads related to size^3, 5, 6^.

While, some obtained data indicate the hair loop and segmental structure of chromosomes in human sperm^7^. Numerous fundamental bio-molecular of sperm have been searched and just a few of these characteristics have been used as tools for medical assessment prior to ART^8, 9^.

More and less analysis of biomarkers such as apoptosis agents, DNA methylation ^9-12^, and measurement of epigenetic subjects ^13, 14^ are not just medical detectors but all demonstrate a piece of the puzzle of sperm structure even sometimes in dinucleotide level ^6, 15^. Some assays such as sperm DNA fragmentation (SDF) have explored chromatin compaction for clinical detection of fertilization outcome^16, 17^.That indicate to the crucial role of condensation in continuing pregnancy^18^ and contrary some reports show there is no correlation between the level of sperm chromatin abnormalities and embryo conception, suggesting more evaluation of chromatin quality^19^. The controversies over the effects of chromatin packaging indicated more research in the sperm nucleus detail.

Different kinds of probes, methods and techniques have been used for diagnostic analysis of sperm in medicine, generating a huge amount of knowledge regarding the structure of sperm. Despite all research, the internal structure and organelles in the sperm head still have great fogginess in chromatids status. Further structural studies conduct better management of diagnostic methods and evaluation of sperm quality in male fertility. For these purposes here the molecular and morphological structure of sperm was studied via quantitative methods and microscopy to approach the micro detail of this cell.

## Methods

### Semen preparation

In this research the available human semen samples were collected for study of the common structure of human sperm. Regardless of the donor’s health or sperm quality the aim of experiments has been access to basic components of sperm structure including chemistry of DNA in different gradients of sperm washes namely 50% and 100% of sperm. These experiments adapted from published dissertation and occurred on semen of a panel of 17 healthy volunteer student donors in the Reproductive Science Research Group at the University of Newcastle, Newcastle, Australia^11^. The base of healthy definition for semen donors had been clinical and para-clinical reports in annual University examination and WHO standards of semen analysis. None of collected semen samples were excluded from the experiments except those that the DNA material were lost through the prolonged process of purification or were low count sperm based on standard semen analysis before and after gradient centrifugation. Through these experiments the semen donors were young in the age range of 20 to 24 and fertility of them had not been a matter of this study. All of the collected samples were gathered after 3 days of sexual abstinence and semen’s were produced by masturbation, collected into sterile containers. Applied method for purification in Newcastle was routine in lab as previously described 50% and 100% discontinuous Percoll (GE Healthcare, Castle Hill, Australia) centrifugation gradients for normal sperm^11^. Use of these samples was approved by the University of Newcastle Human Ethics Committee for a published Mphil dissertation in 2010. And consent was obtained from all participants from the Newcastle University for scientific application of donated samples privately. All methods were performed in accordance with the relevant guidelines and regulations.

### HPLC

HPLC chromatography was applied for evaluation of nucleotides and 5meC in human male germ cells. Sinsheimer’s methods were applied for enzymatic digestion, purification, hydroxylation and quantification components of nucleotides^20^. Extracted and highly purified DNA of human spermatozoa was treated with RNAse Cocktail (Ambion, Austin, TX), following phenol-chloroform purification for 15 minutes prior to injection to HPLC. The mobile phase was 3.51 gr NaH_2_PO_4_ into 960 mL water = 0.03mM and pH adjusted to 5.3 with NH_4_OH. (40mL methanol was added). Then every sample absorbance is monitored at 260 nm and 270 nm instrument and correction factor 5000. Peak areas measurement was quantified with Star Reviewer Software (Varian, Palo Alto, CA). The technique is based on Mossman quantification of the 5-methylcytosine compared with dCMP methylation^21^. The retention time has been expressed near to the certain time since some of picks have occurred with delay because of environment effects on the sequenced elution. The peaks eluted at ∼7 min, ∼15 min, ∼16 min, ∼19min, ∼24min respectively corresponds to 2′-deoxycytidine (227.1 kD), 5-methyl-2′-deoxycytidine (241.2 kD) and three other peaks represent other DNA bases. The area under the peaks of mdCMP and dCMP and other picks did not convert to the molar equivalents by dividing the areas by the extinction coefficients of the respective nucleotides but for quantitation of peak areas we used Star Reviewer software (Varian Inc) result list. This content of bases separated in certain elution times and volume level of picks were measured and listed as percentages in the result column. In order to calibrate the system, deoxyribonucleotide 5m-monophosphate standards dissolved in water were used and the molarities measured according to Beer’s law ^22^.

### Flow cytometry

For the first time flow cytometry has been applied in order to provide an independent assessment of global DNA methylation status in human sperm. After semen sample preparation, 20 × 10^6^ spermatozoa were isolated from each Percoll fraction. For every individual experiment, two controls were used, one unlabeled (without primary or secondary antibody), while the second involved incubation with secondary antibody alone. In the first stage of the analysis spermatozoa were fixed with ethanol (70%) (Merck, Germany) at –20°C for 20 min. Cell pellets were then washed twice in phosphate-buffered saline with Tween 0.5% (Sigma; PBS-T) and centrifuged for 5 min at 500 *g*. The spermatozoa were then incubated at room temperature in 1 Mol/l Tris-HCl buffer, pH 9.5 (Merck), containing 25 mM dithiothreitol (DTT; Sigma) for 20 min after which the samples were washed twice in PBS-T. To ensure that the methylated DNA was accessible to antibody, the sperm chromatin was further denatured with HCl (6 N) for 10 min. This followed washing with Tris-HCl buffer 2 x and once with PBS-T, 1 x. At this stage, spermatozoa for the two control incubations were set aside. The main sample was incubated with anti-5-methylcytosine antibody we employed a protocol similar to that described by Benchaib and his colleague for DNA preparation, then using a mouse monoclonal antibody against 5-methylcytosine (ab73938; Abcam, Sapphire Bioscience, Redfern), diluted 1/100 in PBS-T for 30 min. After two times washes with PBS-T, FITC labeled secondary antibody (anti-mouse IgG), diluted 1/300 in PBS-T was added to the cell pellets and controls containing cells incubated with buffer, instead of the primary antibody. The secondary antibody incubations were conducted for 30 min at 37°C. The cells were then washed twice in PBS-T and run into the flow cytometer. Analyses were performed using the FACS Caliber flow cytometer (BD Biosciences, San Jose, CA). Fluoro chromes were excited with the 488 nm line of the Enterprise laser (Coherent, San Jose, CA). Green fluorescence was detected using FL1 detectors, through band pass filters of 530/30 nm. All data was analyzed with Cell Quest Pro 3.1 software (BD Biosciences).

### CMA3 assay

The spermatozoa recovered from the high and low-density regions of Percoll gradients showed the internal structure and patterns ^23^. Cells were collected (3×10^5^) from each layer, washed and fixed with 100 µl of 4% formaldehyde for 15 minutes. The cell pellets were then washed three times with PBS-T. Then an appropriate aliquot of the cell suspension was placed on a poly-L-Lysine coated coverslip and allowed to settle down overnight at 4°C. Then 50 µl of 0.2% Triton X-100 was added to every coverslip incubated at room temperature for 15 minutes and rinsed with McIlvaine buffer. For CMA_3_ staining, 24 coverslips were incubated with 20 µl of CMA_3_ solution (0.25 mg/ml CMA3= 25 µl of CMA3 stock/mL McIlvaine buffer, pH 7.0) for 20 min. They were then washed in McIlvaine buffer, air-dried and confocal microscopy analysis was then performed using a total of 100 cells per field in every fraction of sperm counted randomly and bright yellow-green cells scored as unprotected positive CMA3 because of protamine deficiency and those stained to dull yellow-green considered as negative CMA3. These were expressed by a bright positive CMA3 percentage cell to the total of 100 stained sperm. The percentage of sperm head with bright green fluorescence was calculated for every slide and the mean of three slides was used for analysis and microscopy process.

### Immunocytochemistry

The method employed for preparing DNA to visualize 5-methyl cytosine residues was essentially that described. To ensure adequate antibody access to its antigen, spermatozoa (20 × 10_5_) were fixed in 100% ice-cold ethanol at -20°C for 15 min; the DNA was then decondensed and permeabilized using HCl and DTT as described above. The slides were rinsed with PBS and blocked for 30 min with 10% goat serum and 1% BSA-PBS. After rinsing with PBS, 5-methylcytosine monoclonal antibody was added in a humidified chamber. Following incubation for 30 min with primary antibody diluted in 1% BSA-PBS at 37°C, the samples were washed three times for 5 minutes with PBS. Finally, the cells incubated for 1h with a secondary antibody, Alexa Fluor goat anti-mouse IgG, diluted in 1% BSA-PBS and washed three times for 5 min with PBS. Washed Cells were mounted with MOWIOL then covered with a coverslip and sealed. Antibody localization was finally visualized with a Zeiss Axioplan 2 fluorescence microscope (Zeiss Filterset 09.^®^ III (Zeiss, Oberkochen, Germany) and confocal microscope.

### Statistical analysis

The HPLC results analyzed by software (Varian Inc). Statistical analyses were performed using Prism version 4.0 (Graphpad software, San Diago, CA, USA). Differences with a probability of *P* < 0.05 were considered statistically significant and calculated by SPSS 18 software.

## Results

## Discussion

### HPLC

In this particular experiment the main concern was measuring the level of nucleotides in sperm. Since hundreds of samples have been lost through the process of preparation, repeated purifications and quantifications, these novel results have culminated of remained12 final samples quantified by HPLC. Through this process the DNA were achieved from low or high percoll gradients and not necessarily paired, from the same individual. Final analysed curves were from 8 students in age ranges of 20 to 24 (Figure 1). The mean area under every typical eluted histogram of micro components in sperm listed in the result column by (Varian Inc) table1. The result was arbitrarily analyzed by percentage of bases in the same time retention was used for calculation of the mean area under peaks. As it showed the mean result of the same peaks from all samples, ranged by sequence from 27.6%, 8.92%, 27.05% and 35.36% (Table. 1). Overall the unique results show that the mean level of major bases, expressed as a percentage of the whole pool of nucleotides. This significant data does not complete the old matched the major nucleotides in human sperm structure.

**Table 1.**
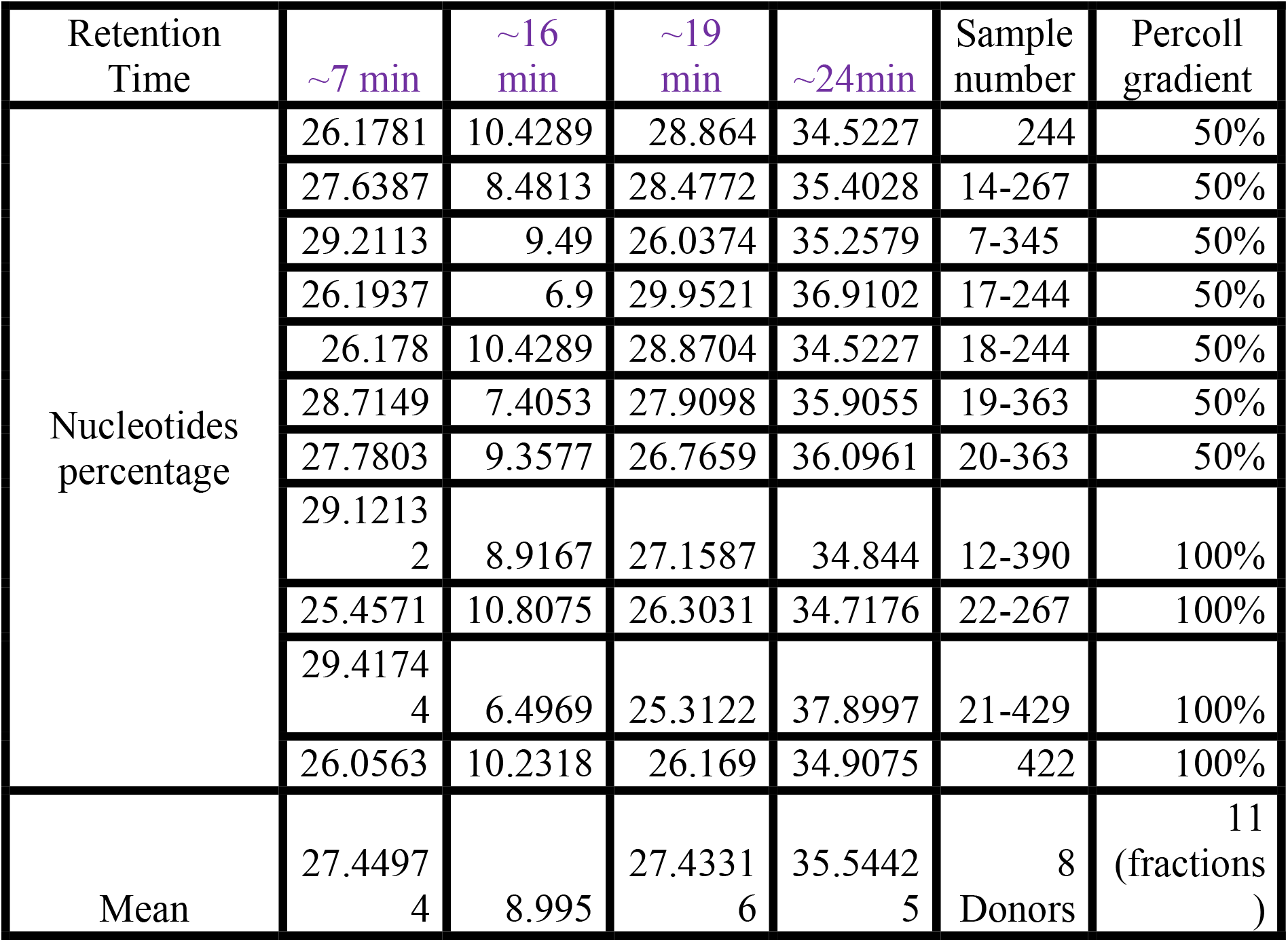

**Figure1.**
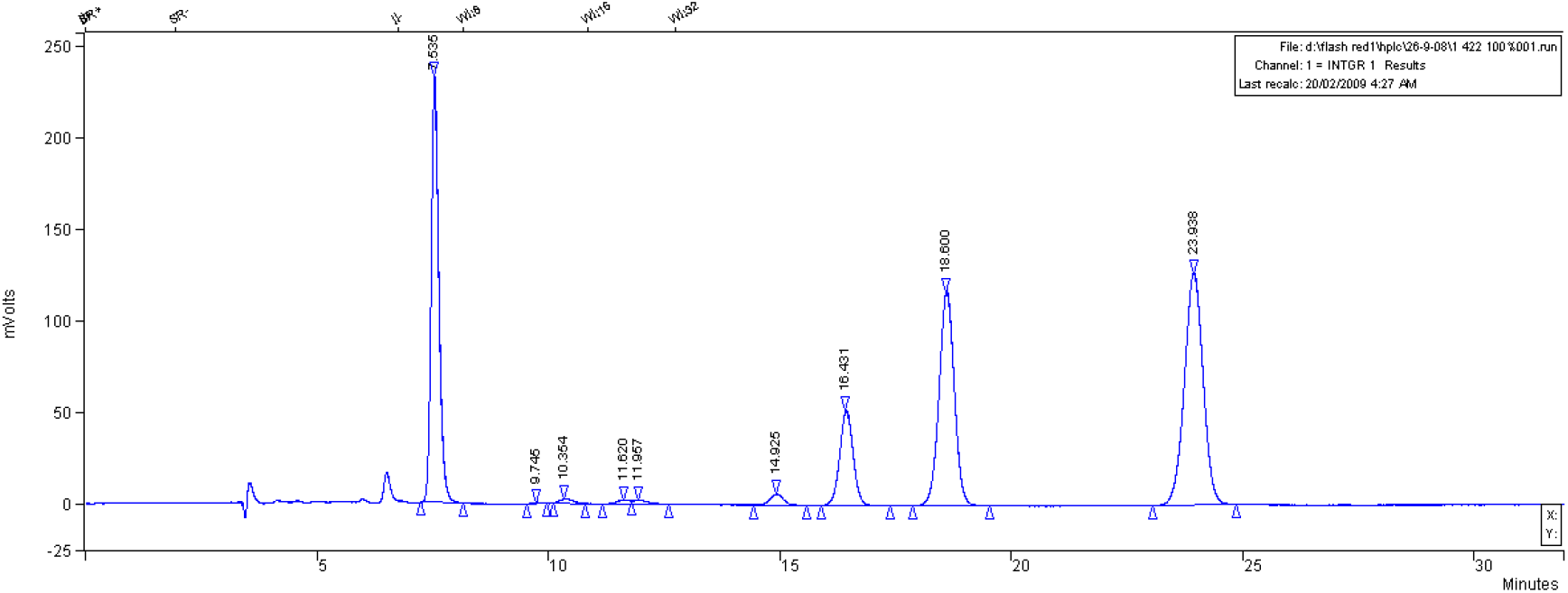
Typical HPLC chromatogram profile of hydrolyzed human sperm DNA and eluted peaks corresponds to bases include 5-m cytosine.

### Flow cytometry

The fluctuation of 5-methylcytosine in sperm DNA structure of every human has proved in this context. This structural element also showed variation in different sperm populations collected from the same sample after percoll gradients wash. The 5-methylcytosine antibody used for 34 percoll washed samples run to the FL2 flow cytometry that showed 100% fraction with the Mean± SEM indicated to14.37±2.719 and expressed levels of Max =56.80 and Min=14.37 in compare to the same levels in gradients of 50% with the Mean± SEM state of 37.00±3.893, Max =92.66 and Min=37.00. In all this study demonstrated that every human male possesses a very individual level of 5-methylcytosine in his sperm (Fig.2).

**Figure 2.**
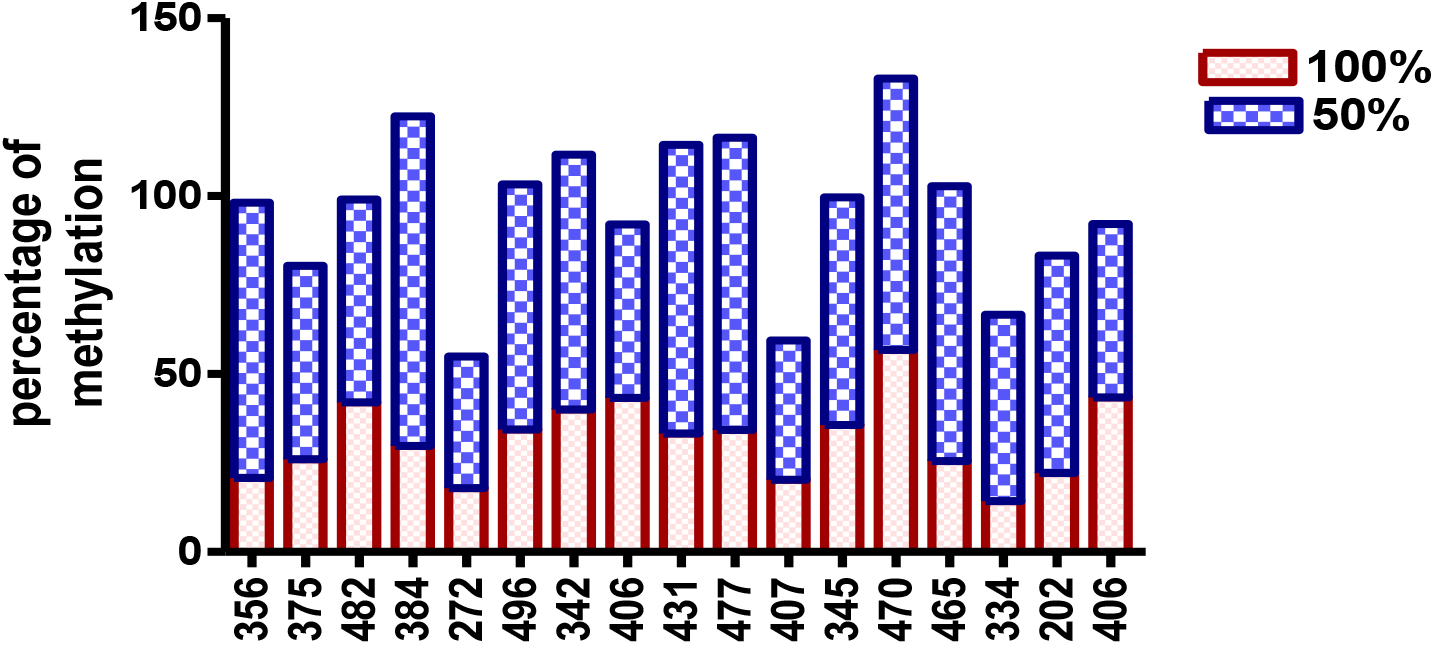
Flowcytometry analysis of 5-methyl cytosine antibody expression from 34 Sperm fractions collected from the high-density region of the gradient (100%) and poor-quality cells (50%) from 17 healthy donors. Every number indicated to a semen donor and pink part of every column demonstrated the level of methylation in50% percoll gradients and blue part pointed to the methylation status in 100% fraction of sperm collected from same individual semen.

### CMA3

The obtained microscopic images of CMA3 staining shows different levels in human spermatozoa recovered from the high and low-density regions of discontinuous Percoll gradients. Comparison of mean expression results of 15 paired samples from the high and low-density regions of percoll gradients were analyzed by SPSS with *P* < 0.0001(Fig.3). Calculation of the percentage of CMA3 positivity stained spermatozoa in 100 sperm per field demonstrated a low level of CMA3 brightness (positivity) in the structure of 100% percoll washed sperm (Fig.5 a-b). Confocal photos of CMA3 stained cells have demonstrated that chromatids are established in a defined spatial geometry through the sperm head (Fig.5 c-d). The image J software has measured the angle between the arm shapes structures presented in sperm DNA by CMA3 staining (Fig.5e).

**Figure3.**
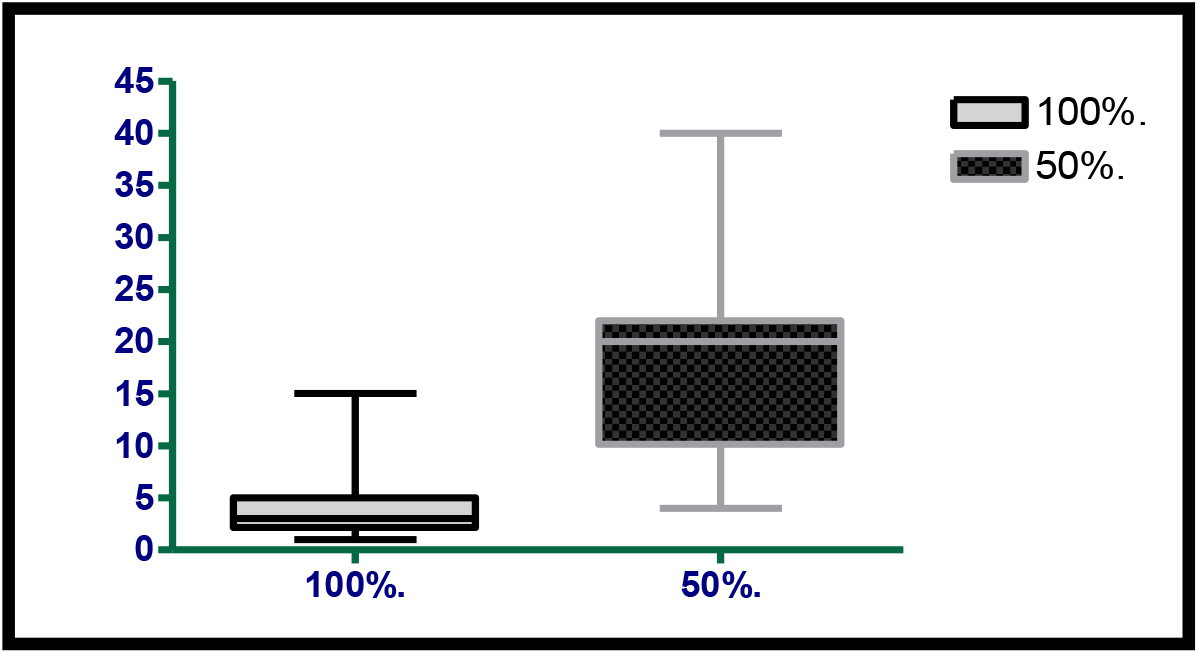
Showing the mean ± SEM level of CMA3 expression in two different populations of spermatozoa prepared on Percoll gradients, analyzed by confocal green fluorescent microscopy. The number of bright cells (CMA3stained) has been expressed as a percentage of cells in a field and compared in the two poor (50%) and fair (100%) populations.

### Immunocytochemistry 5-mC

Immunocytochemical analysis of DNA methylation in the high density Percoll fraction exhibited different patterns of fluorescent signals in comparison to the low density recovered sperm. Microscopic observation demonstrated that the sperms with big heads and cells collected from 50% percoll gradient wash are mostly associated with DNA in hypermethylated stages (Fig.5f-i). Our observation presented a different exhibition with less methylation in high density sperm percoll fraction as it shown in (Fig. j).

### CMA3 and 5-methylcytosine labelled sperm

After Percoll preparation of human sperm, two different fractions were collected and stained with CMA3 and compared with 5-methylcytosine antibody assessment in the same aliquot of relevant fraction measured by flow cytometry. The results show a direct correlation (0.76) between percentage level of CMA3 brightness and the status of DNA methylation from 12 fractions collected from 6 semen donors. This is to compare the overall levels of methylation and chromatin compaction in the same donors analyzed by paired student T-Test with p<0.004 (Fig.4).

**Figure4.**
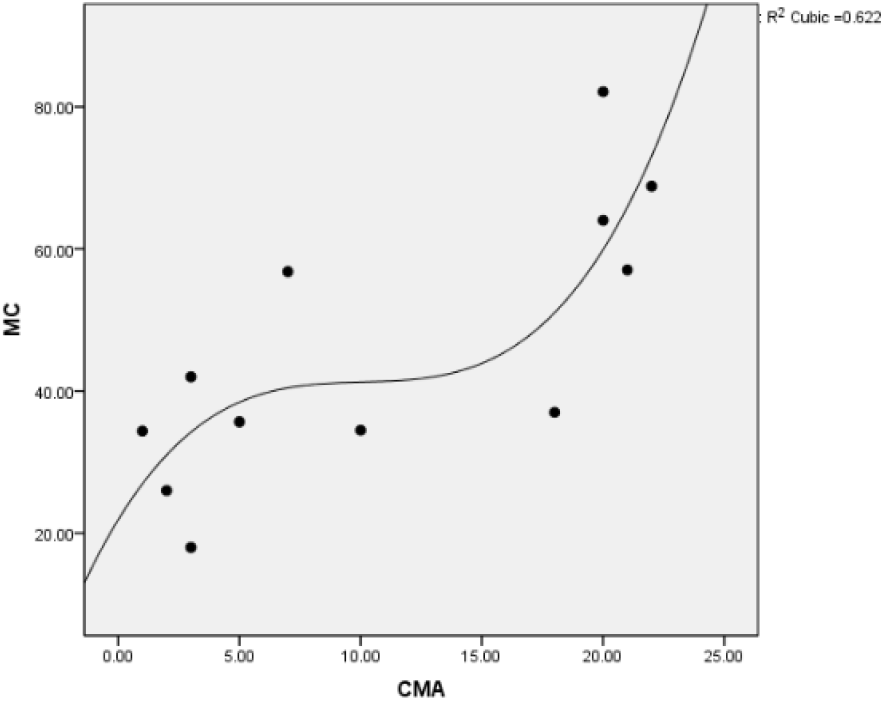
Scatter gram shows the relationship between CMA3 (CMA) staining level and flowcytometery of 5-methyl cytosine (MC) status in 6 semen donors (12 percoll gradient fractions).

### Cytogenetic

Staining occurred on remaining sperms collected from infertility center clients. **Gimsa banding** Sperms samples after swim up wash colored by giemsa banding used for microscopy. Captured photos exhibited by aged and fresh stains indicate a partial model of DNA and morphology of human sperm structure. We used aged and fresh slides for gimsa (G-) banding following trypsin treatment and burned the cell membrane over flame to investigate the pattern of DNA in human spermatozoa. Moreover, the captured photos have demonstrated various aspects of sperm DNA structure that have absorbed the giemsa dye (Fig. p-o).

Our knowledge about DNA alignment, chromatin structure and interpretation of biomarkers are quite limited. Success in the assisted reproductive field currently happens in regard to considering epigenetic markers transmission through generation by the selected best gametes ^26, 27^ and via magnifying the effect of ART on the pool of fertility genes in the future population ^28^. Recent reports have enough attention to treatment of idiopathic infertility with specific methods for prevention of sperm DNA damage ^29^. We have provided first preliminary information to step in producing a whole diagram from the head of sperm that in future made every manipulation based on more visualization. Our findings based on analytical HPLC results for the first time indicate the percentage level of bases in human sperm DNA. We also showed the direct correlation between the 5-methylcytosine levels with CMA3 positivity percentages in sperms collected from low density percoll gradients of these cells. This paper has stepped in peculiar morphology and molecular structure of the sperm galaxy via most advanced and adjusted available methods.

In the present report two quantitative analysis methods were demonstrated the measurements of human sperm components. The samples were carefully processed in extraction, purification, enzymatic digestion, amplification and quantification. However, the uncertainties might occur on the base’s concentration in regards to base’s extinction coefficient, selected path length complications and impurities as well pH and its turbidity that change the light-scattering. For reduction of some of these obstacles we used the concentration of bases without applying the extinction coefficients of four volatiles. Flow Cytometry analysis acknowledges the fluctuation in the achieved results of HPLC, the level of global and individual status of 5-methylcytosine (Fig.2). As it anticipated the mean percentage of two major DNA nucleotides, showing the equal percentage and results does not support the current idea about two by two equality of nucleotides.

Further, CMA3 biomarker compared two different qualities of sperm (Fig.3). CMA3 brightness shows a direct association with the level of 5-methylcytosine (Fig.4), presented in two percoll gradients (Fig.5a-b). A pattern of 5-methylCytosine Corporation has been illustrated (Fig.5f-j). Previously CMA3 brightness was used for checking the DNA quality in human sperm^24^ and karyotyping^30^. The obtained images of sperm nuclear DNA stained by CMA3 (Fig.5c-d) may be pointing to a non-random localization of chromosomes^31^ also is an approbation to evidence for the specific location of individual chromosomes in sperm ^32, 33^. Our finding pointed to CMA3 positivity in some chromatids in cells that are negative to this dye, interpreting that even in normal sperm cells some special chromatids or positions attract CMA3.

**Figure5.**
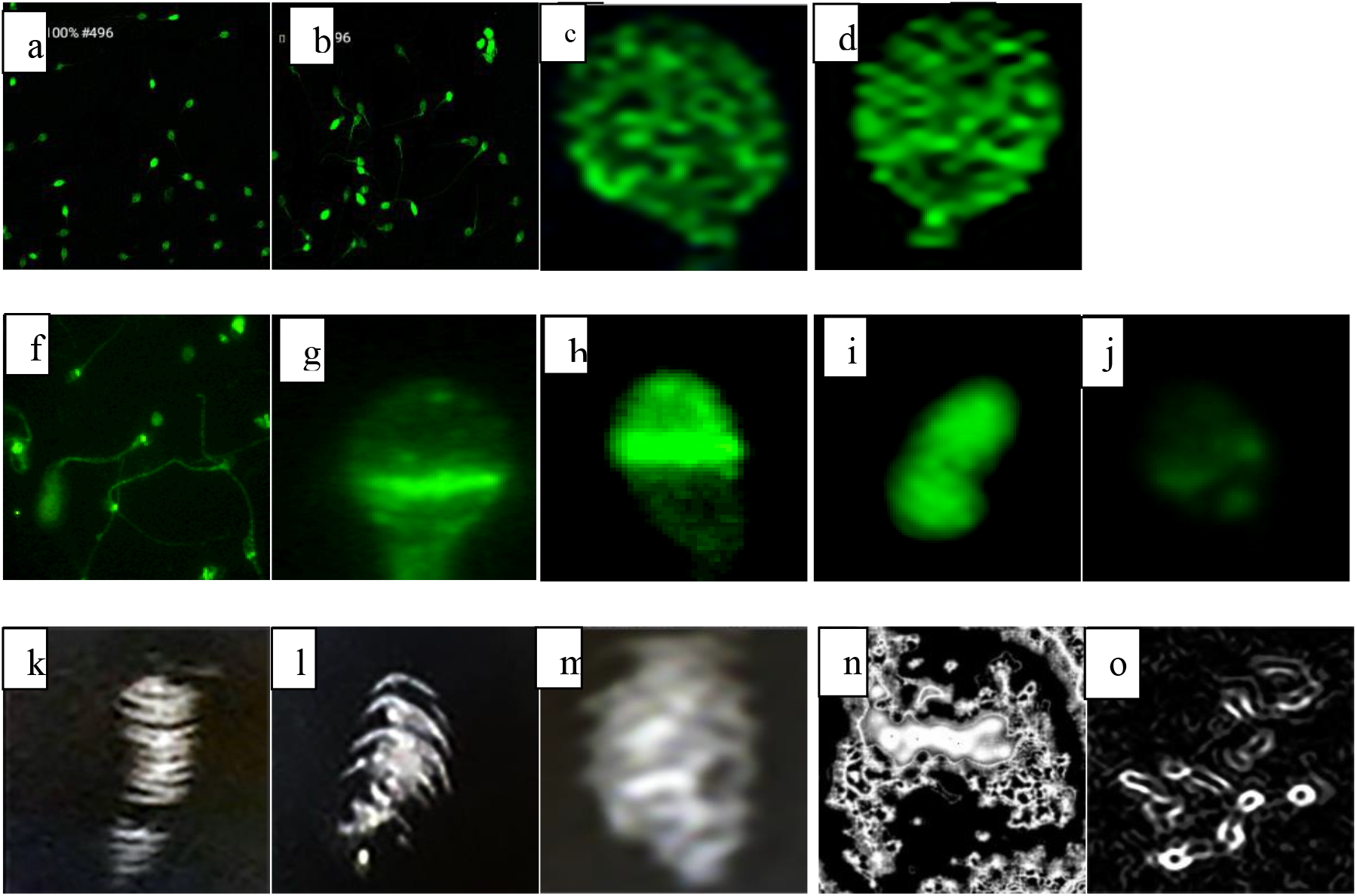
Chromatids’ spatial relation in human sperm. (a-d) CMA3 visualized by green fluorescent confocal microscope. (f-j) pattern of 5-methyl cytosine antibody binding in human sperm. (j) Hypomethylated site observed in fair quality washed. (k-n) geimsa detected by light microscope. (o) geimsa banding pattern was demonstrated very similar to lymphocytes. These micro photos of human sperm structure approve previous reports based on segmental organization of chromatin as well hairpin structure of DNA^7^.

Another particular subject of interest in this study has been providing a 5-methylcytosine pattern in different populations achieved by sperm gradient wash. The photos associated with hypermethylated cells induced the idea of different structure and stages of differentiation in sperm (Fig.5f-i). While the 5-methylcytosine level presented in sperm was lower than the amount expressed in blood DNA^34^ and ovum that indicated to paternal demethylated genes ^35^. Based on evidence, the level of cytosine and 5-m cytosine is not only exclusive to every individual (Fig.2) but to every sperm solely (Fig.5f-j). Through every replacement of –H agent in cytosine by –CH3 the molecular weight increased by fourteen units and ionic charge of DNA changes with addition of two more protons by methyl agent. Future research could uncover the real role of lack of methyl agents in some sperms.

Moreover, captured micrographs of cytogenetic banding giemsa deal with varieties in the sperm demos that partially have been overlooked in the CMA3 and 5-methylcytosine staining methods. The images compromising that every dye exhibit a special part of the human sperm head and variety in the expression of the same antibody might be reflected in structural differences. Analytical quantification and micrograph features are two different views of sperm DNA structure that make a concordance of reviews on dimensional character of chromosomes ^36, 37^. Uncovering the chromatids architecture could have a profound implication on chromatids geometry detection and promising further nuclear manipulation. Through this paper we are compromising other reports that are trying to prove that sperm is more than a silent carrier^38^.

## Conclusion

Further studies based on this research can improve the model of healthy and sick sperm structure but in this text, we are going to step through the first stage and suggest the novel way to enter. Revealing the whole sperm architecture is the first step exploring what is going over the blind technological manipulations. The most important achievement of our study is the listed-out results of Star Varian Inc that urged chemists to follow bases’ percentages equilibrium in axiomatic, namely ladder of human sperm DNA. Moreover, the pattern of 5m-cytosine is exclusive to every individual based on flowcytometry measurement and even to every sperm DNA by micro photography resolution. The microscopy data shed the light on the pieces of logical hologram of the human sperm puzzle map. Our early identification of microscopically and analytical evaluation of male germ line components, need to be completed, support just partial parts of current ideas about sperm DNA. Demands for research in this area are urgent because of human responsibility for consciously managing health and the increase produced by ART.

## List of abbreviations

(ART): Assisted Reproductive Technology
(CMA3): ChromomycinA3
(5-mC): 5-Methylcytosine
(HPLC): High Performance Liquid Chromatography
(dCMP): deoxycytidine monophosphate
(DTT): Dithiothreitol
(BSA): Bovine serum albumin
(PBS): Phosphate-buffered saline

## Authors’ contribution

Not Applicable

## Acknowledgement

Gratefully appreciated prof. Rodney Scott beside his staff Dr David Mossman and Prof. John Aitken and his staff for providing laboratory and supplies in Newcastle University, Newcastle, Australia. Thank to Cyrus Rostami gynecologist and Alireza Qannadi embryologist for providing samples and laboratory facilities from Shiraz Fertility and infertility center, Shiraz, Iran. Also, thank to Dr Fariborz Azad Dr Akbar Safaee for providing pathology facilities in Shiraz University of Medical Sciences, Shiraz, Iran. Thanks to Dr Leila Tahmasbi who admitted me in the training course of karyotyping from Iranian Red Crescent, Tehran, Iran. Final thanks to Gholamreza Hatam, Basic Sciences in Infectious Diseases Research Center, Shiraz University of Medical Sciences, Shiraz, Iran.

